# Assessing Structures and Conformational Ensembles of Apo and Holo Protein States Using Randomized Alanine Sequence Scanning Combined with Shallow Subsampling in AlphaFold2 : Insights and Lessons from Predictions of Functional Allosteric Conformations

**DOI:** 10.1101/2024.11.04.621947

**Authors:** Nishank Raisinghani, Vedant Parikh, Brandon Foley, Gennady Verkhivker

## Abstract

Proteins often exist in multiple conformational states, influenced by the binding of ligands or substrates. The study of these states, particularly the apo (unbound) and holo (ligand-bound) forms, is crucial for understanding protein function, dynamics, and interactions. In the current study, we use AlphaFold2 that combines randomized alanine sequence masking with shallow multiple sequence alignment subsampling to expand the conformational diversity of the predicted structural ensembles and capture conformational changes between apo and holo protein forms. Using several well-established datasets of structurally diverse apo-holo protein pairs, the proposed approach enables robust predictions of apo and holo structures and conformational ensembles, while also displaying notably similar dynamics distributions. These observations are consistent with the view that the intrinsic dynamics of allosteric proteins is defined by the structural topology of the fold and favors conserved conformational motions driven by soft modes. Our findings support the notion that AlphaFold2 approaches can yield reasonable accuracy in predicting minor conformational adjustments between apo and holo states, especially for proteins with moderate localized changes upon ligand binding. However, for large, hinge-like domain movements, AlphaFold2 tends to predict the most stable domain orientation which is typically the apo form rather than the full range of functional conformations characteristic of the holo ensemble. These results indicate that robust modeling of functional protein states may require more accurate characterization of flexible regions in functional conformations and detection of high energy conformations. By incorporating a wider variety of protein structures in training datasets including both apo and holo forms, the model can learn to recognize and predict the structural changes that occur upon ligand binding.

## Introduction

AlphaFold2 (AF2) represents a groundbreaking advancement in the field of protein structure modeling, significantly transforming structural biology [1,2]. AF2 harnesses evolutionary insights from Multiple Sequence Alignments (MSAs) of related protein sequences and employs a hierarchical transformer architecture with self-attention mechanisms, allowing the model to focus on different parts of the protein sequence and understand how distant parts of the sequence interact with each other. AF2’s approach is inspired by natural language processing (NLP) models [3,4], particularly those that use attention-based and transformer mechanisms. These models have revolutionized how computers understand and generate human language, and AF2 applies similar principles to understand the “language” of proteins. By training on vast datasets of protein sequences, AF2 can capture the contextual spatial relationships within proteins, leading to highly accurate structure predictions [1,2]. A recent breakthrough in AI-driven protein structure prediction is represented by ESMFold, an innovative tool developed by Meta AI [5]. ESMFold leverages the ESM2 protein language model (PLM) to predict the three-dimensional structures of proteins directly from their amino acid sequences. This approach eliminates the need for MSAs which are used in AF2 to infer protein structures by comparing sequences of related proteins [5]. These models are trained on vast datasets of protein sequences, enabling them to capture intricate patterns and relationships within the data and can generate highly accurate predictions of protein structures at the atomic level, significantly speeding up the process compared to AF2 methods. OmegaFold is another powerful method that Predicts high-resolution protein structures directly from a single primary sequence without invoking MSAs [6]. This method uses a combination of PLM and a geometry-inspired transformer model. Although AF2-based methods and self-supervised PLM approaches have made significant strides in predicting static protein structures, they face notable limitations in their ability to characterize conformational dynamics, functional protein ensembles, conformational changes, and allosteric states [7]. Recent studies suggest that although AF2 methods excel at predicting individual protein structures, they encounter significant challenges in accurately modeling conformational ensembles and mapping allosteric landscapes [8–12]. This issue may arise from a training bias towards experimentally validated, thermodynamically stable structures, and MSAs that primarily capture evolutionary information aimed at predicting ground states. The recent study applied AF2 methods for detecting both apo and holo states of 91 proteins showing that AF2 performance worsens with the increasing conformational diversity of the studied protein systems [13]. Although AF2 methods showed robust performance in predicting the experimentally determined ground conformation for 98 fold switching proteins, they typically failed to detect alternative structures suggesting the inherent AF2 network bias for the most probable conformer rather than an ensemble of relevant functional states [14]. The revealed biases limit the ability of AF2 methods to capture the diversity of protein conformational dynamics [15–18] and expanding AF2’s predictive range to include functional conformational ensembles and reliably map allosteric landscapes is now a central focus of rapidly growing computational research in this area [19–25].

Recent adaptations to the AF2 framework that target the prediction of alternative protein conformational states employ innovative strategies, such as reducing the depth of the MSA by sampling only a subset of sequences, thereby creating a shallower MSA [19]. This strategy aims to enhance sequence diversity within the model, potentially broadening its capacity to capture a range of alternative conformational states. Another approach, SPEACH_AF (Sampling Protein Ensembles and Conformational Heterogeneity with AlphaFold2), uses in silico alanine mutagenesis within MSAs to expand AF2’s attention network, enabling a deeper exploration of coevolutionary residue patterns associated with different conformations [20]. This technique leverages the dynamic interactions of residues, enhancing AF2’s ability to map conformational heterogeneity and identify functionally relevant alternative states. Together, these adaptations offer promising pathways for advancing AF2 beyond single-structure predictions, potentially facilitating more accurate modeling of protein flexibility and allosteric mechanisms. Structure prediction dep leaning network Cfold, which is trained on a conformational split of the Protein Data Bank (PDB) to generate alternative conformations, enables efficient exploration of the conformational landscape of monomeric protein structures with over 50% of experimentally known nonredundant alternative protein conformations predicted with high accuracy [21]. Clustering MSAs by sequence similarity allows AF2 to sample alternative states of known metamorphic proteins with high confidence [22]. This method, termed AF-Cluster, allows predictions of alternative protein states and has demonstrated success in identifying previously unknown fold-switched states, which were subsequently validated using NMR analysis [22]. The latest analysis of the AF2 methods tested the predictive ability on fold-switching proteins, showing relatively weak reproducibility of experimental fold switching, failure to discriminate between low and high energy conformations and suggesting a strong bias towards ground states arising from structural memorization of training-set structures rather than from understanding of protein thermodynamics [23]. Recent AF2 adaptations integrate sequence and evolutionary data derived from MSAs, as well as structural insights obtained from templates. This approach has been particularly successful in applications to proteins like kinases and GPCRs [24]. AF2 methodologies were also actively explored for predicting conformational states in protein kinases. AF2-based modeling of 437 human protein kinases in the active form using shallow MSAs of orthologs and close homologs of the query protein showed the robustness of AF2 methods as selected models for each kinase based on the prediction confidence scores of the activation loop residues conformed closely to the substrate-bound experimental structures [25]. The ability of AF2 methods to predict kinase structures in different conformations at various MSA depths was examined, demonstrating that using lower MSA depths allows for more efficient exploration of alternative kinase conformations [26]. By exploring different adaptations of the MSA subsampling architecture, a latest insightful study systematically examined the ability of the AF2 method to characterize the conformational distributions of the ABL kinase domain and predict the effects of state-switching allosteric mutants [27]. Computational studies showed the intrinsic difficulties and limitations of conventional biophysical simulations to accurately characterize transient states and describe kinetics of conformational changes as even extremely long MD simulations coupled with the enhanced sampling approaches often fail to detect functionally relevant alternative conformations of protein kinases and unable to accurately map complex allosteric landscapes [28].We recently proposed a new AF2 adaptation in which randomized alanine sequence scanning (AF2-RASS) of the entire protein sequence or specific functional regions is combined with the MSA subsampling enabling interpretable atomistic predictions and adequate characterization of the ABL conformational ensembles for the active and inactive states [29,30]. These studies suggested that key challenges of the emerging AF2 adaptations are associated with accurate predictions of functional conformations and relative populations of distinct allosteric states rather than simply increasing the conformational diversity of the predicted ensembles. Current developments highlight key challenges in AF2 methodologies and their adaptations. These include accurately capturing functional conformational ensembles, understanding allosteric states, and elucidating the impacts of mutations on both local and global protein conformational changes.

Proteins often exist in multiple conformational states and prediction of the apo (unbound) and holo (ligand-bound) forms, is crucial for understanding protein function, dynamics, and interactions. A computational approach to predict the structure of protein/ligand complexes based solely on the unbound conformation and the ligand data was initially proposed and tested on apo-holo proteins that undergo substantial structural rearrangements upon binding [31]. Protein flexibility upon ligand binding was analyzed using 305 proteins with 2369 holo structures and 1679 apo structures that were obtained using Binding MOAD[32] followed by filtering for proteins with at least 2 holo structures and 2 apo structures [33]. This study showed that apo and holo structures can exhibit similar structural variation as measured by residual backbone RMSD changes. A nonredundant dataset of 521 paired protein structures in the apo and holo forms was used to estimate the degree of both local and global structure similarity between the apo and holo forms, showing that most apo-holo protein pairs did not exhibit a significant structural difference [34]. Another approach generated reliable holo-like structures from apo structures by ligand binding site refinement with restraints derived from holo templates with low homology demonstrating the apo structures can be refined toward the corresponding holo conformations for 23 of 32 proteins of the DUD-E data set [35]. Several computational databases have been developed to facilitate the study of apo-holo pairs. Apo–Holo DataBase (AH-DB), contains 746 314 apo–holo structure pairs of 3638 proteins from 702 organisms [36]. Apo-Holo Juxtaposition (AHoJ) web application was developed for retrieving apo-holo structure pairs for user-defined ligands [37]. AHoJ-DB (www.apoholo.cz/db) database was built by matching the binding sites of biologically relevant protein–ligand interactions from the PDB with their apo and holo counterparts [38]. Conformational Diversity in the Native State of proteins (CoDNaS) database represents another source of apo and holo conformational samples [39,40]. AF2 approaches, while highly effective at predicting single, static structures of proteins, have limitations when it comes to distinguishing between different conformational states, such as apo (unbound) and holo (ligand-bound) forms. AF2 models generally predict the most thermodynamically stable, ground-state structure, which often resembles either an apo form or a general conformation that might not fully capture ligand-induced structural changes. Recent AF2 adaptations and strategies, such as AF2 with MSA subsampling or using clustering methods such as AF-Cluster [22] can expand AF2’s prediction capability to capture structural variations closer to those seen in different functional states, including apo-holo pairs. Nonetheless, achieving accurate apo and holo predictions remains a challenge, as AF2 [1,2], AlphaFold-multimer [41] or AF2Complex [42] are trained on experimentally determined, stable structures without specific emphasis on conformational dynamics or allosteric changes associated with ligand binding. AF2 is reasonably effective at predicting small ligand-induced adjustments, especially for binding sites with minor side-chain rearrangements or small-scale domain shifts. However, larger-scale movements, such as domain swapping or substantial loop reorganization upon ligand binding, are often beyond its predictive capacity without further adaptation. A recent study applied AF2 methods for detecting both apo and holo states of 91 proteins showing that for 67% of the proteins the AF2 models had the lowest RMSD to the holo form, while only 33% of the proteins are modeled with lowest RMSD to the apo form [13]. The key finding of this study showed that in a curated collection of apo–holo pairs of conformers, AF2 is unable to reproduce the observed conformational diversity with the same error for both apo and holo conformers. Overall, AF2’s capabilities have expanded our understanding of static protein structure but continue to face challenges in accurately modeling the dynamic, functional states of proteins, including detailed apo-holo transitions and allosteric mechanisms.

In the current study, we use AF2-RASS adaptation to explore conformational changes between apo and holo protein forms. Using several well-established datasets of structurally diverse apo-holo protein pairs, we predicted structures and conformational ensembles of the unbound and bound protein states. The results of this study demonstrate that unlike standard AF2 approach, AF2-RASS adaptation can adequately sample both apo and holo protein forms and capture invariant dynamics signatures of the conformational ensembles. We find that combining alanine sequence masking with shallow MSA subsampling can significantly expand the conformational diversity of the predicted structural ensembles and detect alternative protein conformations. We argue that the AF2 RASS adaptation with systematic perturbation of the MSAs through iterative random scanning of the protein sequence can loosen coevolutionary constraints and reduce structural “memorization” allowing for conformational sampling of alternative states.

## Results and Discussion

### Structural Analysis of Datasets of Apo and Holo Protein Pairs Reveals Conformational Variability and Different Motions Induced by Ligand Binding

In the dataset we included 10 apo-holo protein pairs from [31] and 43 pairs from the PocketMiner dataset [43], which is a collection of apo-holo protein structure pairs with significant conformational changes upon ligand binding (Supporting Information, Table S1). The subset of 10 protein pairs includes domain-induced closures upon ligand binding and significant structural rearrangements. In the Pocket Miner dataset, structural changes between apo and holo forms include loop motions in dihydrofolate reductase (apo PDB: 2W9T, holo PDB: 2W9S) [44]; secondary structure movements in lipoprotein LpqN (apo PDB: 6E5D, holo PDB: 6E5F) [45] and integrin-binding protein 1 (apo PDB: 1Y1A chain A, holo PDB: 1Y1A chain B) [46]; and the interdomain motions in nopaline-binding periplasmic protein (apo PDB: 4P0I, holo PDB: 5OTA) [47]. We illustrated the structural differences between apo and holo forms by presenting the aligned conformations of apo and holo protein forms for 12 representative pairs (Figure 1). Significant structural differences between the apo and holo protein forms can be seen for D-Allose binding protein (apo PDB :1gud, holo PDB: 1rpj, RMSD=3.65 Å); D-Ribose binding protein (apo PDB: 1urp; holo PDB :2dri, RMSD = 3.25 Å); 5-Enolpyruvylshikimate-3-phosphate synthase (apo PDB: 1rf5; holo PDB :1rf4, RMSD= 2.99 Å) and osmo-protection protein (apo PDB: 1sw5; holo PDB :1sw4, RMSD= 3.67 Å, TM-score =0.75). (Figure 1C-F). In these cases, we observe large and distinct movements and rearrangements in the holo protein forms. In particular, we can observe hinge-bending motion of D-allose binding protein that exists in a dynamic equilibrium of closed and open conformations where in the closed ligand-bound form D-allose is buried at the domain interface (Figure 1C) [48]. Conformational changes are necessary for the function of bacterial periplasmic receptors in chemotaxis and transport. The open, ligand-free forms of the *Escherichia coli* ribose-binding protein were observed in X-ray crystallographic studies and together with the previously described closed, ligand-bound forms showed that the open forms are related to the closed form by a hinge motion [49]. At the same time, there is a significant number of apo-holo pairs in which ligand binding can induce only moderate local structural variations within RMSD= 1.0-2.0 Å (Figure 1)

**Figure 1.**
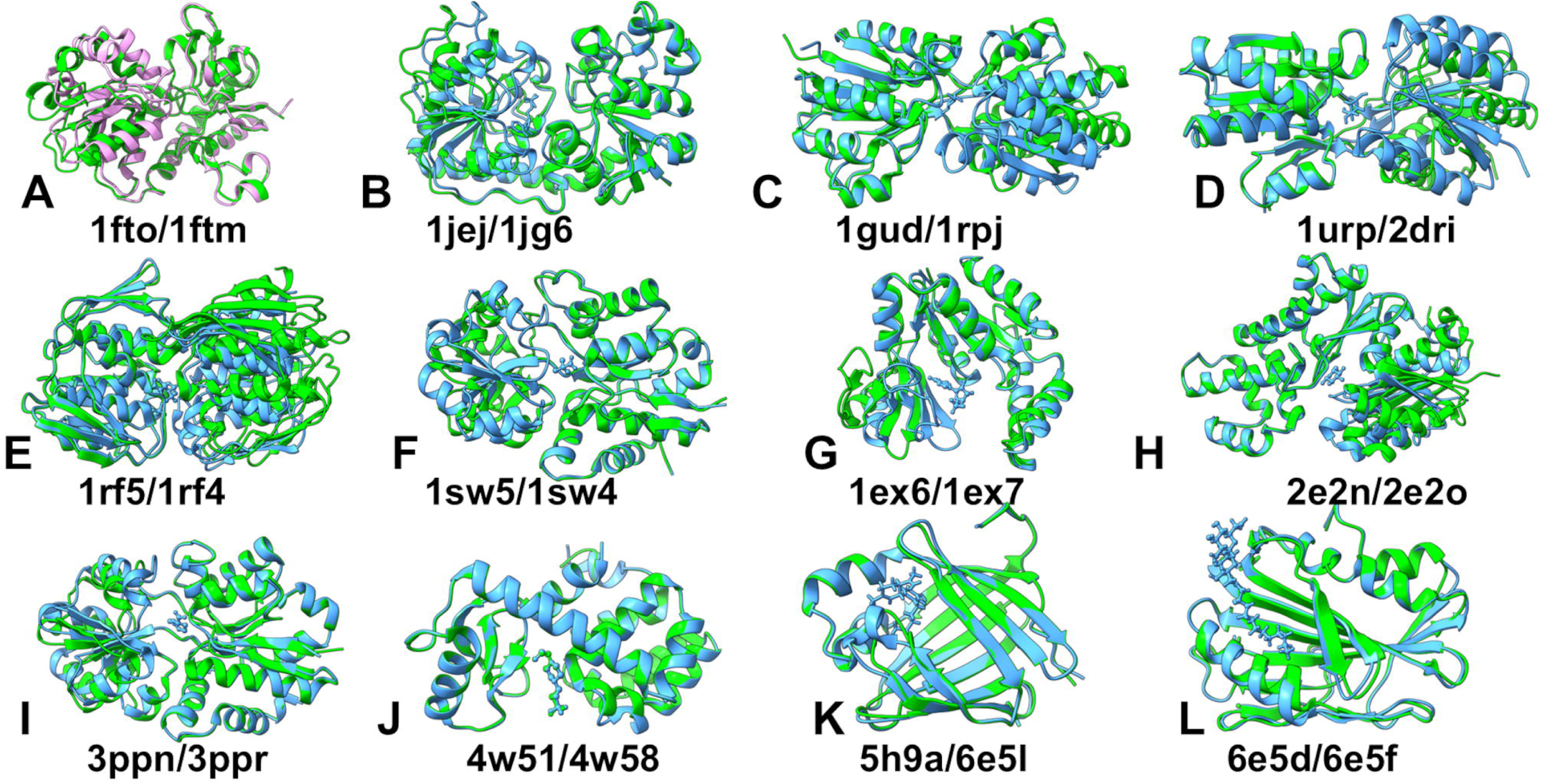
Structural alignment of the representative subset of 12 apo/holo pairs. Apo x-ray structures are shown in green and holo structures are colored in light blue. (A) The apo-holo structure of GluR2 ligand binding core (apo PDB :1fto, holo PDB: 1ftm, RMSD=2.2 Å, TM-score=0.89). (B) DNA Beta-Glucosyl-transferase (apo PDB :1jej, holo PDB: 1jg6, RMSD=2.1 Å, TM-score=0.93). (C) D-Allose binding protein (apo PDB :1gud, holo PDB: 1rpj, RMSD=3.65 Å, TM-scor3=0.76). (D) D-Ribose binding protein (apo PDB: 1urp; holo PDB :2dri, RMSD = 3.25 Å, TM-score=0.77). (E) 5-Enolpyruvylshikimate-3-phosphate synthase (apo PDB: 1rf5; holo PDB :1rf4, RMSD= 2.99 Å, TM-score =0.84). (F) Osmo-protection protein (apo PDB: 1sw5; holo PDB :1sw4, RMSD= 3.67 Å, TM-score =0.75). (G) Guanylate kinase (apo PDB: 1ex6; holo PDB :1ex7, RMSD= 2.98 Å, TM-score =0.82). (H) Hexokinase (apo PDB: 2e2n; holo PDB :2e2o, RMSD= 2.95 Å, TM-score =0.85). (I) ABC transporter OpuC (apo PDB: 3ppn; holo PDB :3ppr, RMSD= 1.46 Å, TM-score =0.94). (J) T4 Lysozyme L99A (apo PDB: 4w51; holo PDB :4w58, RMSD= 0.7 Å, TM-score =0.99). (K) Human cellular retinol binding protein 1 (apo PDB: 5h9a; holo PDB :6e5l, RMSD= 0.99 Å, TM-score =0.97). (L) Lipoprotein LpqN (apo PDB: 6epd; holo PDB :6e5f, RMSD= 0.79 Å, TM-score =0.99).

We computed analyzed the distribution of the RMSD values and TM-score values between apo and holo protein forms in the dataset (Figure 2). The distribution of RMSD values showed at least three different peaks with the largest peak RMSD ∼ 0.7-1.0 Å and other notable peaks at RMSD ∼ 3.0 Å and RMSD ∼ 4.0 Å (Figure 2A). This indicates that the dataset features different levels of conformational changes ranging from structurally similar apo-holo pairs that displayed only small local changes to apo-holo pairs in which ligand binding can induce significant conformational changes and deviations from the unbound protein (Figure 2A). The distribution of TM-score values reflected this trend showing that in addition to major peaks at TM-score ∼ 0.95 there is a broad range of smaller peaks at TM-score ∼ 0.65-0.9. The latter peaks correspond to the apo-holo pairs that featured significant structural changes in the bound protein form (Figure 2B)

**Figure 2.**
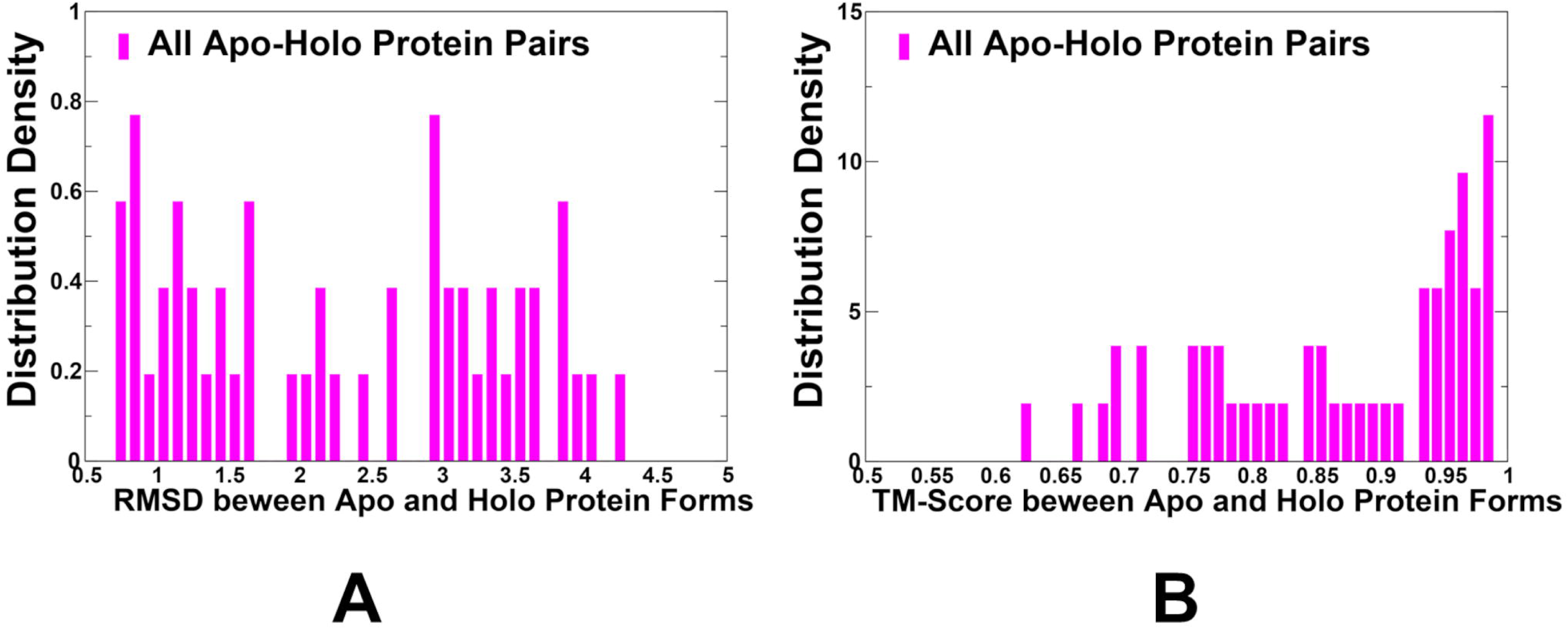
The analysis of the RMSD values and TM-score values between apo and holo protein forms in the dataset of apo-holo protein forms. (A) The density distributions of the RMSD values between apo and respective holo conformations in the dataset. (B) The distribution density of TM-scores estimating structural differences between apo and holo conformations.

### AF2-RASS Adaptation Can Capture Structures and Conformational Ensembles of Apo and Holo Protein Forms

We employed recently developed randomized alanine scanning adaptation of the AF2 methodology in which the algorithm operates first on the pool of sequences and iterates through each amino acid in the native sequence to randomly substitute residues with alanine, thus emulating random alanine mutagenesis [30]. In the proposed protocol, randomized alanine sequence scanning is performed for the entire protein sequence or specific kinase regions involved in conformational changes followed by construction of corresponding MSAs and then by AF2 shallow subsampling applied on each of these MSAs [30]. AF2 estimates prediction quality with two confidence metrics: the per residue predicted Local Difference Distance Test (pLDDT) and predicted template modeling (pTM) scores [1,2]. AF2 models were ranked by pLDDT scores (a per-residue estimate of the prediction confidence on a scale from 0 to 100), quantified by the fraction of predicted Cα distances that lie within their expected intervals. The values correspond to the model’s predicted scores based on the lDDT-Cα metric which is a local superposition-free metric that assesses the atomic displacements of the residues in the predicted model. To gain a quantitative insight into the AF2-RASS predictions, we constructed the pLDDT density distribution for the predicted conformational ensembles of the (Figure 3). The dominant peaks at pLDDT ∼90 and shallow peaks for pLDDT∼75-80 are indicative of high quality conformations in the predicted ensembles (Figure 3). By reranking the predicted conformations according to the percentage of confident residues we selected the stable conformations where the large fraction of residues ( > 70%) featured the high confidence values pLDDT ∼ 85-90 and therefore are assumed to be functionally relevant stable states [50,51]. The predicted models are compared to the experimental structure using structural alignment tool TM-align [52,53]. We used TM-score which is a metric for assessing the topological similarity of protein structures based on their given residue equivalency. TM-score ranges from 0 to 1, where a value of 1 indicates a perfect match between the predicted model and the reference structure. When TM-score > 0.5 implies that the structures share roughly the same fold. TM-score > 0.5 is often used as a threshold to determine if the predicted model has a fold similar to the reference structure.

**Figure 3.**
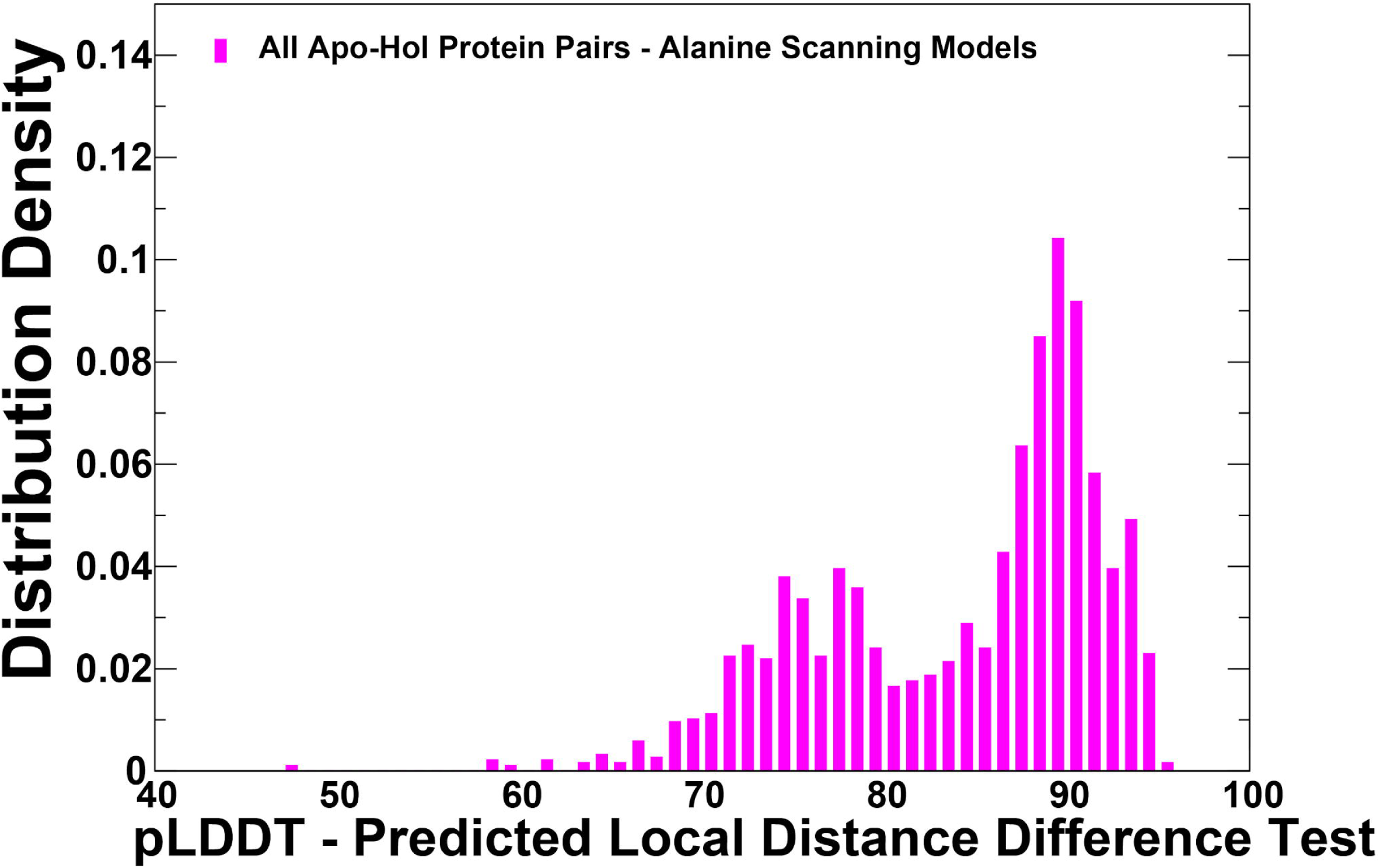
The analysis of AF2-RASS predictions of the conformational ensembles for apo-holo protein forms. The density distributions of the pLDDT estimates for AF2-RASS predicted conformations. The pLDDT structural model estimate of the prediction confidence is on scale from 0 to 100.

The predicted conformational ensembles reflected differences in conformational mobility between protein systems in which apo and holo forms are structurally similar and apo-holo pairs with significant movements in the holo forms. As may be noticed, there is considerable degree of conformational fluctuations in the D-Allose binding protein (Figure 4C), D-Ribose binding protein (Figure 4D), 5-Enolpyruvylshikimate-3-phosphate synthase (Figure 4E) and Osmo-protection protein (Figure 4F). These ensembles displayed both local mobility and reflected on larger structural changes. For other apo-holo pairs, for example T4 Lysozyme L99A (Figure 4J) and Lipoprotein LpqN (Figure 4L) that featured structurally similar apo and holo forms, the observed conformational mobility is largely confined to peripheral regions and flexible loops.

**Figure 4.**
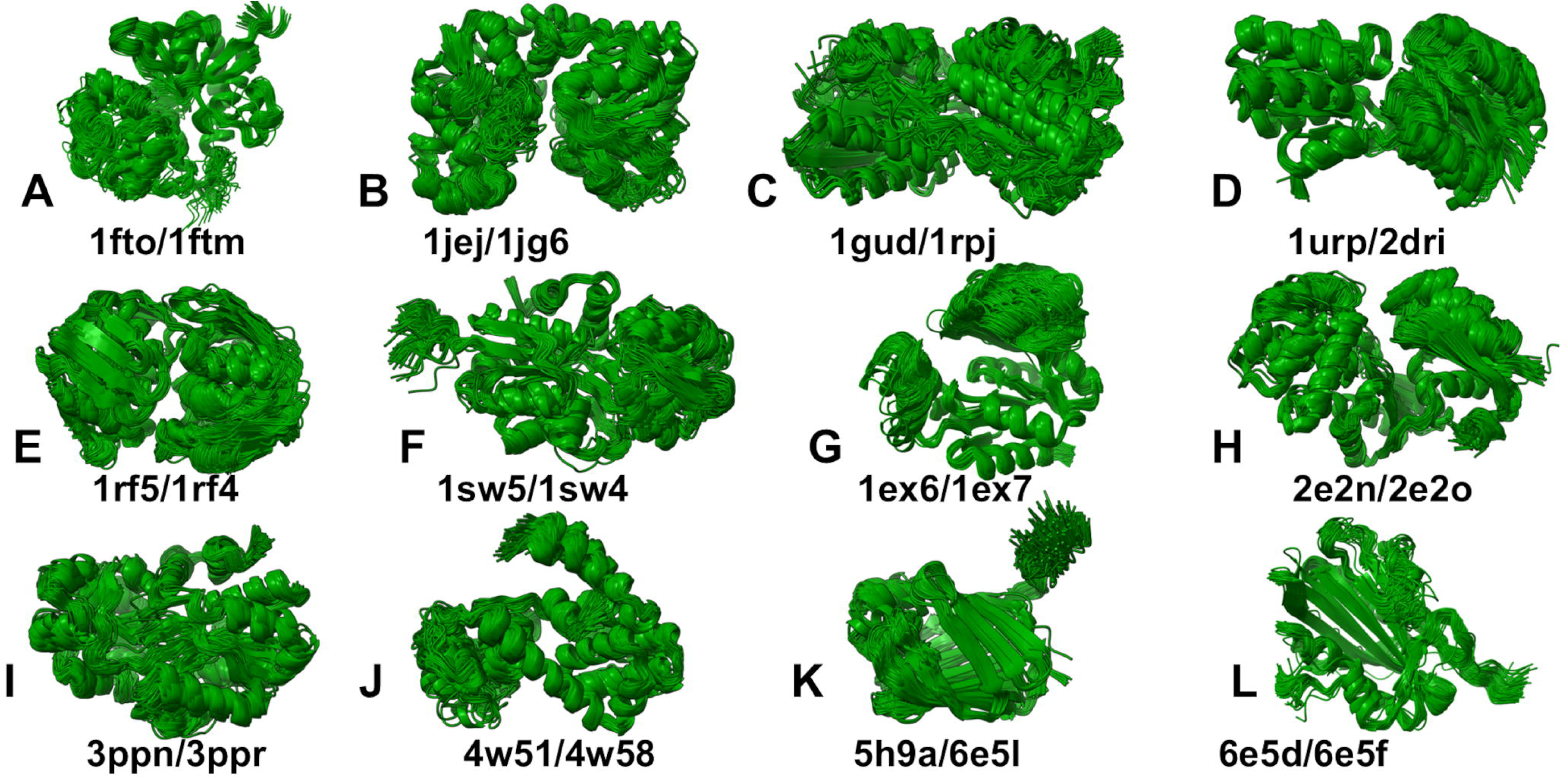
The AF2-RASS predicted conformational ensembles for apo-holo proteins (A) The apo-holo structure of GluR2 ligand binding core (apo PDB :1fto, holo PDB: 1ftm). (B) DNA Beta-Glucosyl-transferase (apo PDB :1jej, holo PDB: 1jg6). (C) D-Allose binding protein (apo PDB :1gud, holo PDB: 1rpj). (D) D-Ribose binding protein (apo PDB: 1urp; holo PDB :2dri). (E) 5-Enolpyruvylshikimate-3-phosphate synthase (apo PDB: 1rf5; holo PDB :1rf4). (F) Osmo-protection protein (apo PDB: 1sw5; holo PDB :1sw4). (G) Guanylate kinase (apo PDB: 1ex6; holo PDB :1ex7). (H) Hexokinase (apo PDB: 2e2n; holo PDB :2e2o). (I) ABC transporter OpuC (apo PDB: 3ppn; holo PDB :3ppr). (J) T4 Lysozyme L99A (apo PDB: 4w51; holo PDB :4w58). (K) Human cellular retinol binding protein 1 (apo PDB: 5h9a; holo PDB :6e5l). (L) Lipoprotein LpqN (apo PDB: 6epd; holo PDB :6e5f).

### Lessons from Modeling of Conformational Ensembles of Apo and Holo Protein Forms : Capturing Shared Signature Dynamics and Ligand-Induced Conformational Changes

We analyzed the global RMSD distributions of the AF2-RASS predicted ensembles with respect to the corresponding apo and holo forms (Figure 5). The shape of the RMSD distributions is similar with pronounced peaks at RMSD ∼1.0-2.0 Å from the crystal structures (Figure 5). The AF2-RASS predicted ensembles demonstrated robust coverage of both apo and holo structural forms, while the explored conformational dynamics reveals similar distributions with respect to apo and holo forms (Figure 5). Our findings are consistent with the notion that a defining characteristic of allosteric proteins is their ability to access highly conserved, global motion modes that are present and shared between apo and holo forms [54]. It was also suggested that functional fitness of proteins is determined by conserved global dynamics of a versatile fold, while gaining specificity of interactions may be achieved via localized fluctuations conserved among subfamily members but divergent across subfamilies [54]. Hence, the obtained conformational ensembles may provide adequate description of main thermal fluctuations taken place in both apo and holo forms.

**Figure 5.**
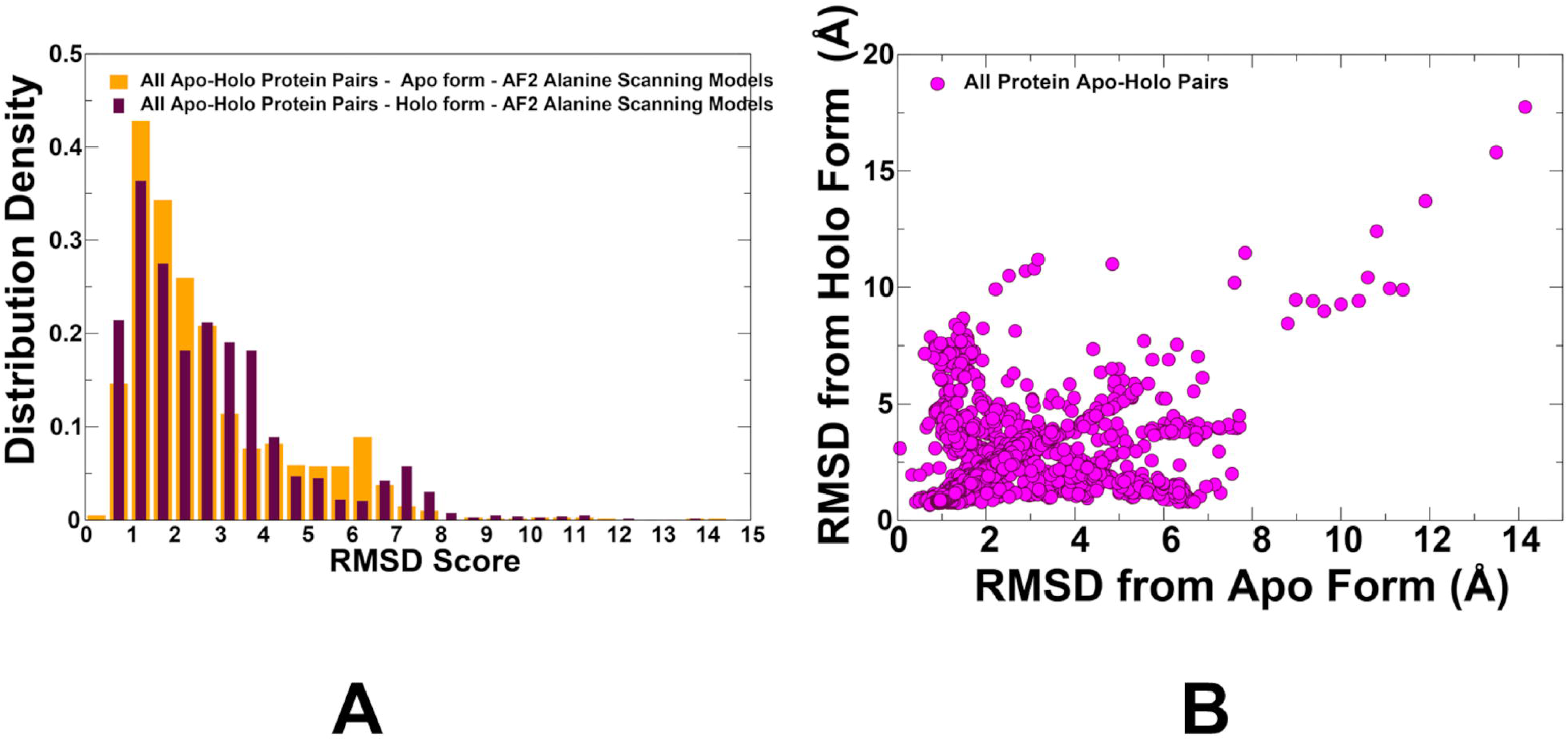
(A) The distribution of the RMSD values between AF2-RASS predicted conformations in the ensemble with respect to the apo crystal structure (in orange bars) and holo structure (in maroon bars). (B) The scatter plot between the RMSD values with respect to the apo and holo structures (shown in magenta-colored filled circles).

We also plotted the distributions of the RMSD values of the conformational ensembles against the apo and holo forms (Figure 6). The average RMSD against their apo form is only slightly lower (mean = 2.14 Å) than against their holo form (mean = 2.39 Å) which highlights the fact that the AF2-RASS predicted ensembles can adequately sample the basins of both apo and holo forms. The box plots also illustrate remarkably similar population distributions for RMSDs with respect to apo and holo forms. This contrasts with previous studies when using AF2 default method it was shown that most of the proteins are modeled with a bias toward a given conformer.

**Figure 6.**
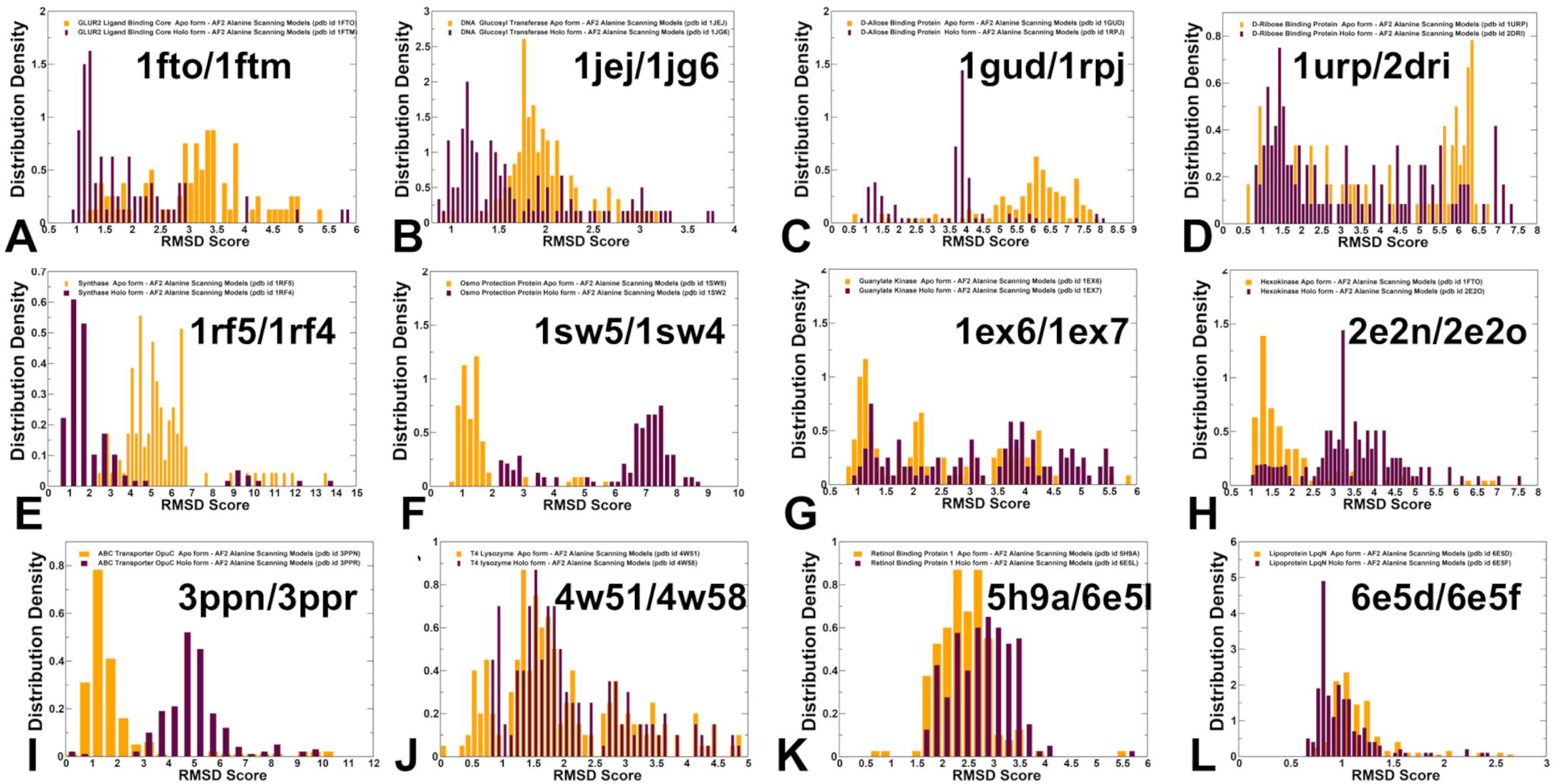
The box plots of the distribution of the RMSD values between AF2-RASS predicted conformations in the ensemble with respect to the apo crystal structure (A) and holo structure (B).

In order to understand better the nature of the conformational ensembles for apo-holo protein forms and differences between AF2-RASS sampling of these states for studied systems, we analyzed the distributions of the RMSD values between AF2-RASS predicted conformations in the ensemble with respect to the apo and holo crystal structures (Figure 7). Importantly, we found that for the vast majority of studied apo-holo pairs, AF2-RASS can produce conformational ensembles that sample both apo and holo structures. However, in a few cases of osmo-protection protein (apo PDB: 1sw5; holo PDB :1sw4) and ABC transporter OpuC (apo PDB: 3ppn; holo PDB :3ppr) the generated ensemble is biased towards the native apo conformation, while the holo structure could not be adequately sampled in simulations (Figure 7).

**Figure 7.**
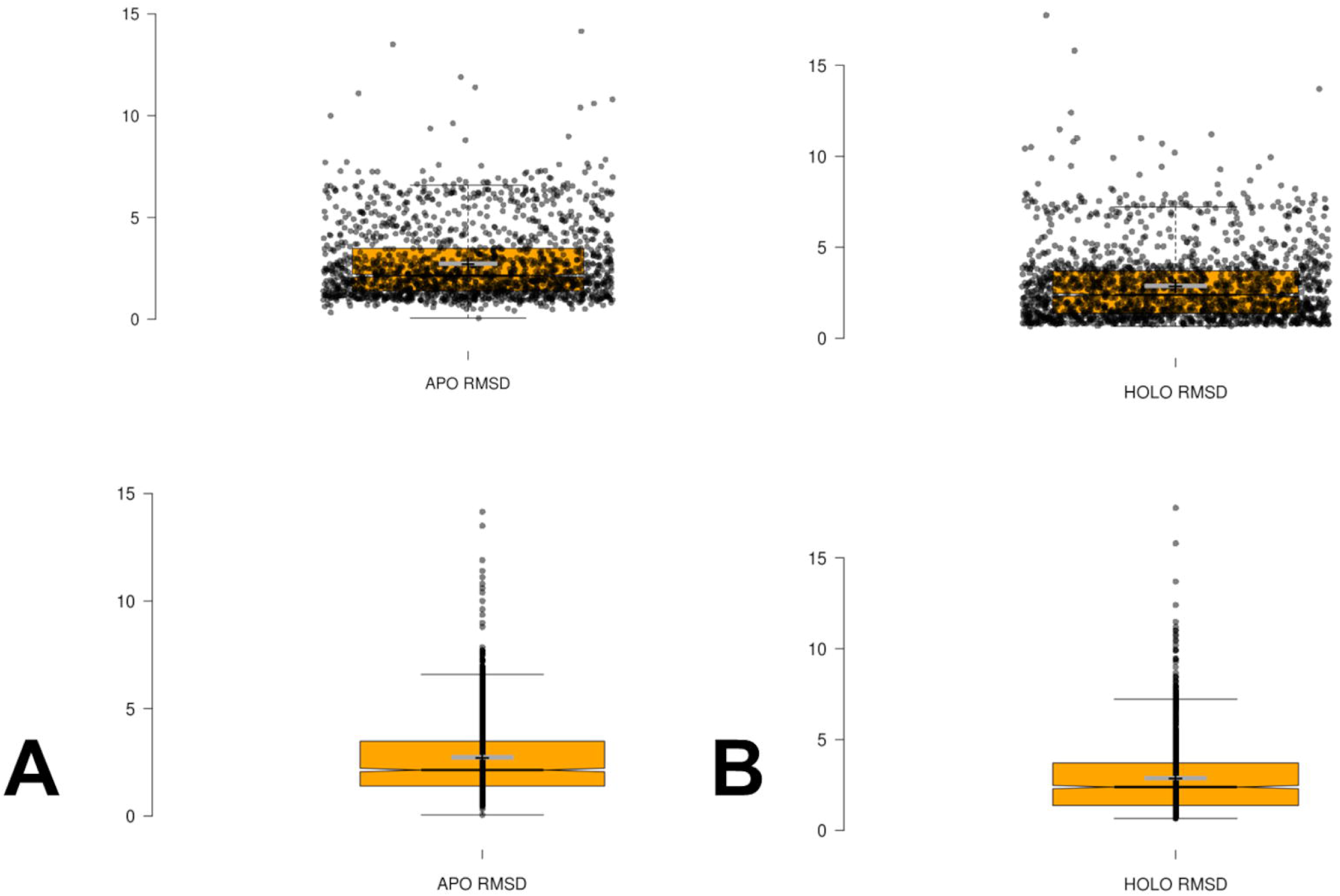
The distribution of the RMSD values between AF2-RASS predicted conformations in the ensemble with respect to the apo crystal structure (in orange bars) and holo structure (in maroon bars). (A) The apo-holo structure of GluR2 ligand binding core (apo PDB :1fto, holo PDB: 1ftm). (B) DNA Beta-Glucosyl-transferase (apo PDB :1jej, holo PDB: 1jg6). (C) D-Allose binding protein (apo PDB :1gud, holo PDB: 1rpj). (D) D-Ribose binding protein (apo PDB: 1urp; holo PDB :2dri). (E) 5-Enolpyruvylshikimate-3-phosphate synthase (apo PDB: 1rf5; holo PDB :1rf4). (F) Osmo-protectid on protein (apo PDB: 1sw5; holo PDB :1sw4). (G) Guanylate kinase (apo PDB: 1ex6; holo PDB :1ex7). (H) Hexokinase (apo PDB: 2e2n; holo PDB :2e2o). (I) ABC transporter OpuC (apo PDB: 3ppn; holo PDB :3ppr). (J) T4 Lysozyme L99A (apo PDB: 4w51; holo PDB :4w58). (K) Human cellular retinol binding protein 1 (apo PDB: 5h9a; holo PDB :6e5l). (L) Lipoprotein LpqN (apo PDB: 6epd; holo PDB :6e5f).

The structural analysis of the apo and holo forms of the ligand-binding protein ProX (Figure 1F, 4F) showed significant conformational changes and hinge movements at the inter-domain regions where residues provided by domain A remain at their apo positions, while the protein residues of domain undergo a large conformational shift [55]. According to these structural experiments, the equilibrium between the open and closed forms is shifted toward the closed ligand-bound form. Our results indicated that AF2 approaches including AF2-RASS adaptation, which is designed to enhance sampling of multiple functional states, could exhibit biases towards the native apo form, particularly in cases of unique conformational changes where movements of one of the domain are coupled to ligand binding. A similar mechanism of large and system-specific conformational changes was seen for the substrate-binding protein OpuCC of the ABC transporter OpuC that can recognize a broad spectrum of compatible solutes. Structural studies determined crystal structures of OpuCC in the apo-form and in complex with various ligands [56]. The structures showed that OpuCC is composed of two α/β/α globular sandwich domains linked by two hinge regions, with a substrate-binding pocket located at the interdomain cleft (Figure 1I, 4I). Upon substrate binding, the two domains shift towards each other to trap the substrate where a flexible pocket can accommodate various compatible ligands [56]. Hence, for this apo-holo protein pair, ligand binding can induce large interdomain shifts that are also accompanied by synchronous local conformational adjustments in the binding pocket. Our results showed that AF2-RASS predicted conformations mostly correspond to the apo crystal structure and the generated ensemble consists of conformations in the local vicinity of the apo structure, likely reflecting local conformational fluctuations around this dominant state. These observations pointed to an important fact that performance of AF2 methods in predicting functional conformational diversity and ensembles may depend on the complexity and system-specific nature of conformational changes. We argue that AF2 methods may have learned how to predict structures of the native proteins through structural memorization of databases, which may also enable robust prediction of local conformational changes and even larger changes that are well-represented in the training datasets. However, these methods may fail to recognize more unique conformational rearrangements in holo structures that are driven by non-trivial combination of inter-domain and local changes.

The results showed a strong correlation between conformational flexibility and pLDDT metric for apo-holo pairs in which ligand binding induced local moderate conformational changes within RMSD=2.0-2.5 Å between apo and holo forms (Figure 8).

**Figure 8.**
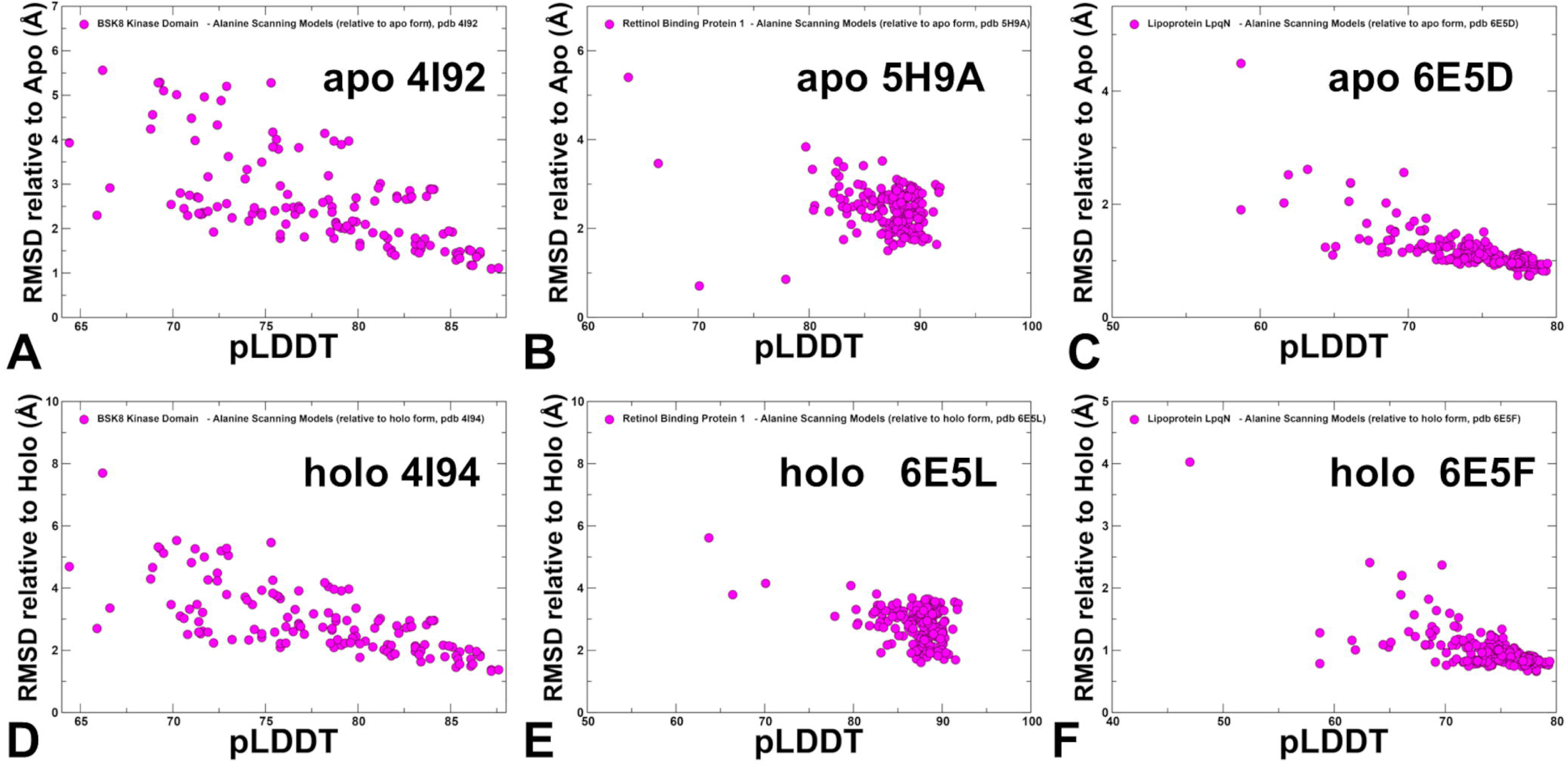
The scatter plots of pLDDT and RMSDs for the conformational ensembles obtained with AF2-RASS approach. The scatter plots between pLDDT and RMSDs from apo crystal structures are shown for BSK8 apo crystal structure , pdb id 4I92 (A), Human cellular retinol binding protein 1 apo structure, pdb id 5H9A (B) and Lipoprotein LpqN apo structure, pdb id 6E5D (C). The scatter plots between pLDDT and RMSDs from holo crystal structures are shown for BSK8 apo crystal structure , pdb id 4I94 (D), Human cellular retinol binding protein 1 holo structure, pdb id 6E5L (E) and Lipoprotein LpqN holo structure, pdb id 6E5F (F). In contrast, for apo-holo pairs exhibiting larger structural changes including loop motions and flexibility changes, the pLDDT-RMSD relationship becomes more complex, reflecting the ruggedness of conformational landscapes and insufficient accuracy in predicting flexible regions involved in modulation of apo-holo transitions (Figure 9).

**Figure 9.**
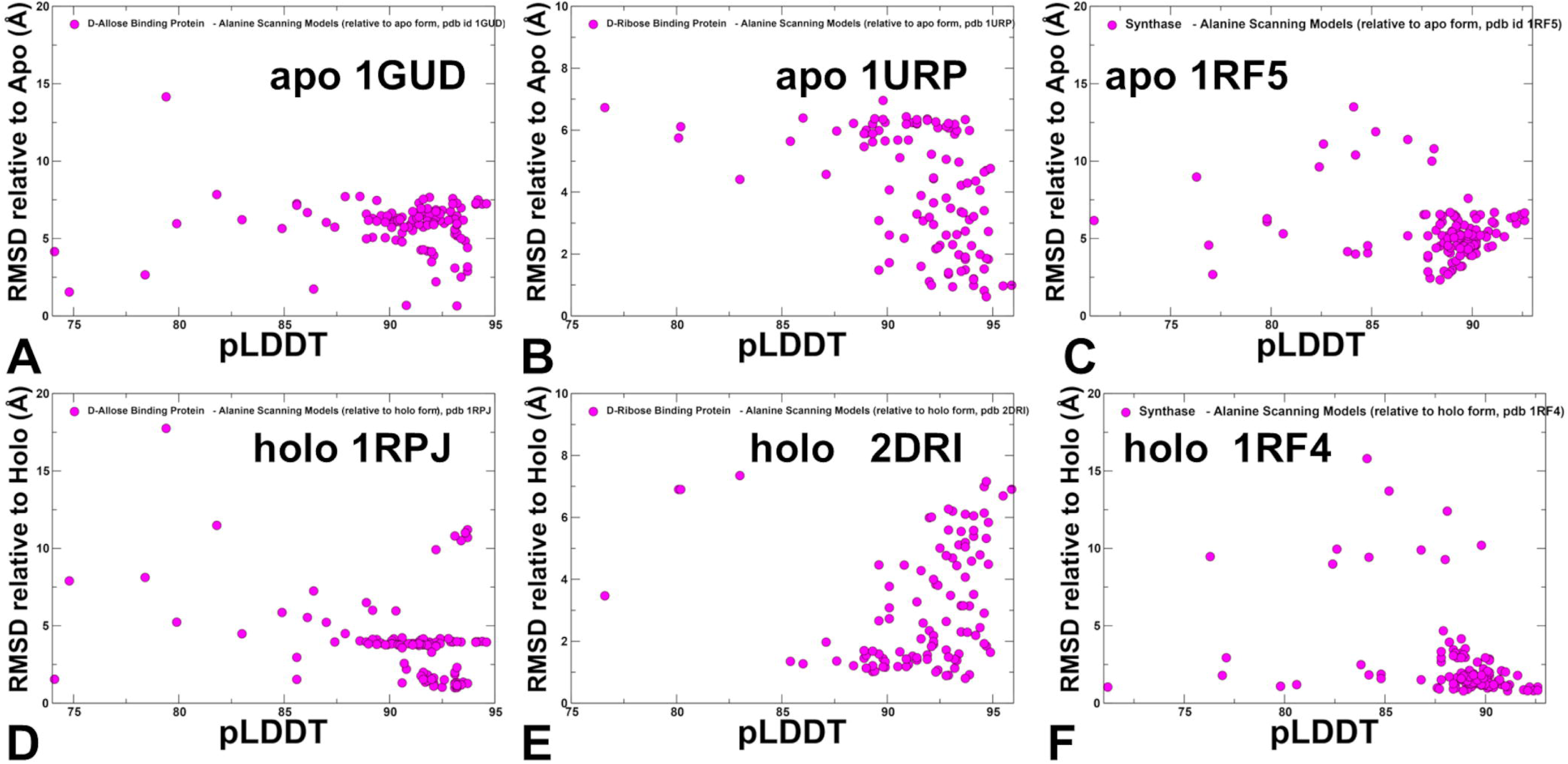
The scatter plots of pLDDT with RMSDs for the conformational ensembles obtained with AF2-RASS approach. The scatter plots between pLDDT and RMSDs from apo crystal structures are shown for D-Allose binding protein apo, pdb id 1GUD (A), D-Ribose binding protein apo, pdb id 1URP (B) and 5-Enolpyruvylshikimate-3-phosphate synthase apo (pdb id 1RF5). The scatter plots between pLDDT and RMSDs from holo crystal structures are shown for D-Allose binding protein holo, pdb id 1RPJ (D), D-Ribose binding protein holo, pdb is 2DRI (E) and 5-Enolpyruvylshikimate-3-phosphate synthase holo, pdb id 1RF (F).

**Figure 10.**
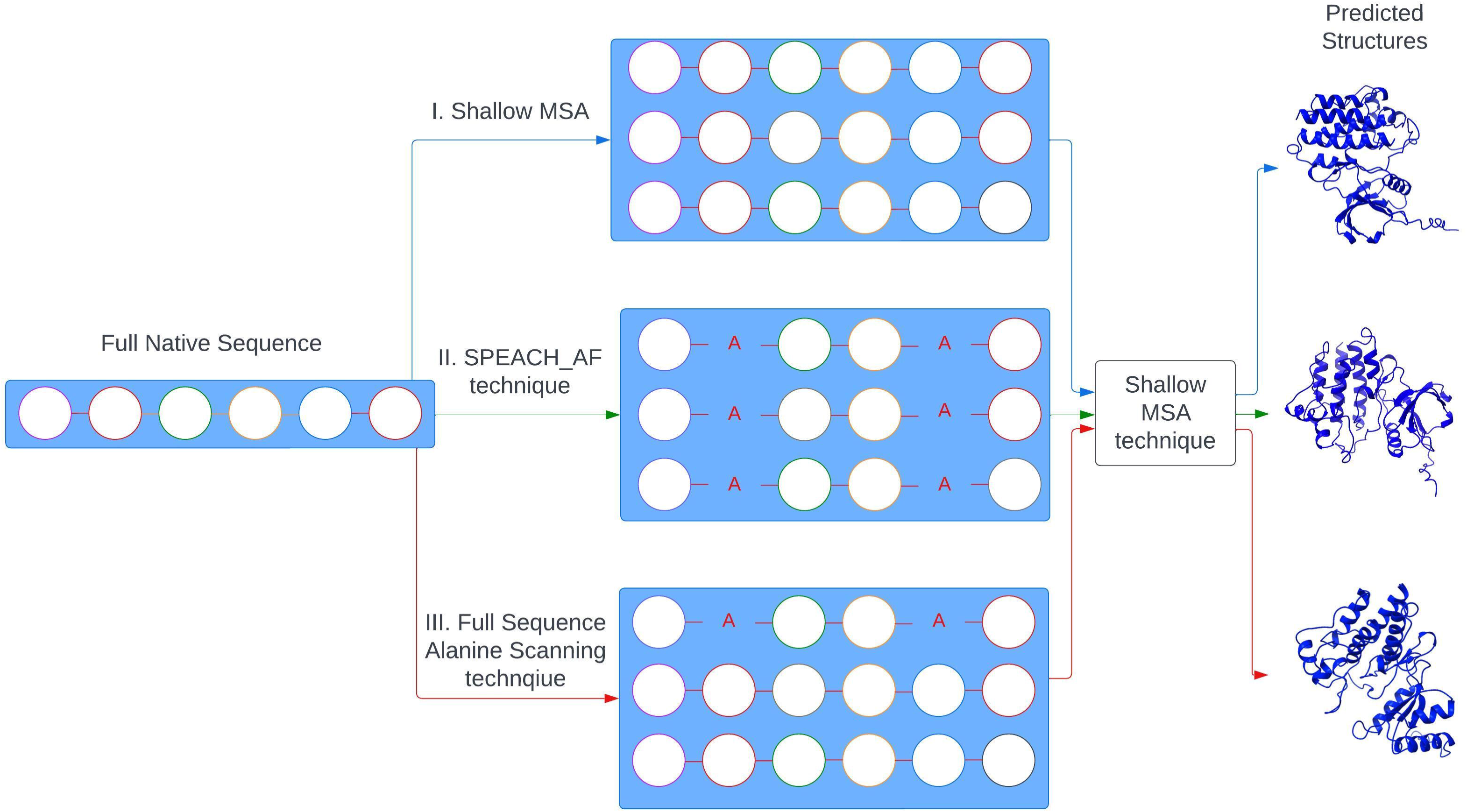
A schematic representation of the AF2 protein structure prediction pipeline using shallow MSA subsampling, SPEACH_AF method and AF2-RASS approach.

The results are consistent with previous studies by Porter and colleagues [23] showing that AF2 confidence metrics can often select against alternative conformations failing to predict the most energetically favorable fold-switch conformations and discriminate between low and high energy conformations. Our findings support the notion that AF2 approaches can yield reasonable accuracy in predicting minor conformational adjustments between apo and holo states, especially for proteins with small, localized changes upon ligand binding such as side-chain reorientations and local flexibility of loop regions. In proteins where ligand binding induces minor domain shifts or rotations, AF2 can predict conformations close to the holo state, provided the structural differences are localized. However, for large, hinge-like domain movements, AF2 tends to predict the most stable domain orientation which is typically the apo form rather than the full range of functional conformations characteristic of the holo ensemble. These results indicate that modeling of multiple functional states of proteins may require more accurate detection of flexible region conformations and cannot solely rely on the pLDDT metric as the major determinant of the prediction accuracy in reproducing functional conformational ensembles.

## Materials and Methods

### AF2 with MSA Shallow Subsampling

Structural predictions were carried out using AF2 framework [1,2] within the ColabFold implementation [57] using a range of MSA depths and MSA subsampling. The MSAs were generated using the MMSeqs2 library [58,59] using the ABL1 sequence from residues 240 to 440 as input. We used *max_msa* field to set two AF2 parameters in the following format: *max_seqs:extra_seqs*. These parameters determine the number of sequences subsampled from the MSA (*max_seqs* sets the number of sequences passed to the row/column attention track and *extra_seqs* the number of sequences additionally processed by the main evoformer stack). The default MSAs are subsampled randomly to obtain shallow MSAs containing as few as five sequences. This parameter is in the format of *max_seqs:extra_seqs* which decides the number of sequences subsampled from the MSA. *Max_seq* determines the number of sequences passed to the row/column attention matrix at the front end of the AF2 architecture, and *extra_seqs* sets the number of extra sequences processed by the Evoformer stack after the attention mechanism. The lower values encourage more diverse predictions but increase the number of misfolded models. We explored the following parameters: *max_seq, extra_seq*, number of seeds, and number of recycles. We ran simulations with *max_seqs:extra_seqs* 16:32, 32:64, 64:128. 128:256, 256:512 and 512:1024 values and report the results at *max_seqs:extra_seqs* 16:32 that produced the greatest diversity. We additionally manipulated the *num_recycles* parameters to produce more diverse outputs. To generate more data, we set *num_recycles* to 12, which produces 14 structures starting from recycle 0 to recycle 12 and generating a final refined structure. Recycling is an iterative refinement process, with each recycled structure getting more precise. AF2 makes predictions using 5 models pretrained with different parameters, and consequently with different weights. Each of these models generates 14 structures, amounting to 70 structures in total. We then set the *num_seed* parameter to 1. This parameter quantifies the number of random seeds to iterate through, ranging from random_seed to random_seed+num_seed. We also enabled the use_dropout parameter, meaning that dropout layers in the model would be active during the time of predictions.

### AF2 with Randomized Alanine Sequence Scanning and Shallow Subsampling of MSAs

The initial input for the full sequence randomized alanine scanning is the original full native sequence. This technique utilizes an algorithm that iterates through each amino acid in the native sequence and randomly substitutes 5-15% of the residues with alanine [30]. The algorithm substitutes residue with alanine at each position with a probability randomly generated between 0.05 and 0.15 for each sequence position. We ran this algorithm multiple times (∼40-50) on the full sequences for each mutant, resulting in a multitude of distinct sequences, each with different frequency and position of alanine mutations. The AF2 shallow MSA methodology is subsequently employed on these MSAs to predict protein structures as described previously. A total of 70 predicted structures were generated from 12 recycles per model.

The root mean square deviation (RMSD) superposition of backbone atoms were calculated using ProFit (http://www.bioinf.org.uk/software/profit/).

## Conclusions

In the current study, we use AF2-RASS adaptation to predict structures and conformational ensembles of apo and holo protein forms using several datasets of structurally diverse apo-holo protein pairs. We found that combing alanine sequence masking with shallow MSA subsampling can significantly expand the conformational diversity of the predicted structural ensembles and detect populations of both apo and holo forms. The AF2-RASS approach enables predictions of both apo and holo structural forms, while also displaying notably similar dynamics distributions. Combining alanine sequence masking with shallow MSA subsampling can significantly expand the conformational diversity of the predicted structural ensembles and detect alternative protein conformations. We argue that the AF2 RASS adaptation with systematic perturbation of the MSAs through iterative random scanning of the protein sequence can loosen coevolutionary constraints and reduce structural “memorization” allowing for conformational sampling of alternative states. Our results highlighted several critical limitations of current models, showing that while AF2 adaptations can accurately predict moderate ligand-induced structural changes between apo and holo forms and produce functionally significant conformational ensembles for unbound and ligand-bound protein states, these approaches may be challenged by large functional transitions involving the inter-domain rearrangements and subtle combination of global and local structural changes. These results indicate that robust modeling of functional protein states may require more accurate characterization of flexible regions in functional conformations and detection of high energy conformations. Identifying and modeling high-energy conformations that proteins might transiently adopt could provide insights into the pathways of conformational changes is currently an unexplored area of AF2 methodologies. Integrating AF2-RASS and other AF2 adaptations with enhanced sampling methods can help capture the physics-based dynamic nature of proteins [60,61]. Developing generative models like AlphaFlow and ESMFlow, which use flow-based generative modeling, may further improve sampling the conformational landscapes of protein and detection of apo and holo structural ensembles [62] . By incorporating a wider variety of protein structures, including both apo and holo forms, the model can learn to recognize and predict the structural changes that occur upon ligand binding. With a richer dataset, the model can refine its predictions, leading to more accurate and reliable structural models. Advances in sampling methods, hybrid models, and adaptive frameworks continue to improve the predictive landscape, opening doors to more detailed and functional protein modeling.

## Supplementary Materials

The following supporting information can be downloaded at: www.mdpi.com/xxx/s1, Table S1: The dataset of apo and holo protein pairs used in this study.

## Data Availability Statement

Data is fully contained within the article and Supplementary Information material. Crystal structures were obtained from Protein Data Bank (http://www.rcsb.org). The rendering of protein structures was done with UCSF ChimeraX package (https://www.rbvi.ucsf.edu/chimerax/) and Pymol (https://pymol.org/2/). The software tools used in this study are available at GitHub sites : https://github.com/deepmind/alphafold; https://github.com/sokrypton/ColabFold/;https://github.com/RSvan/SPEACH_AF; https://www.github.com/HWaymentSteele/AFCluster; https://github.com/nickraisgit/ABL1KinasePaper

All the data obtained in this work, the software tools, and the in-house scripts are freely available at ZENODO open repository: https://zenodo.org/records/14031452.

## Acknowledgments

G.V acknowledges support from Schmid College of Science and Technology at Chapman University for providing computing resources at the Keck Center for Science and Engineering.

